# Dabigatran prevents lipopolysaccharide mediated apoptosis in zebrafish through a thrombin independent mechanism

**DOI:** 10.1101/2025.10.03.680206

**Authors:** Annelore I. T. Fleischmann, William H. Woodhams, Ritta Mouayed, Timmy Joseph, Xiangyu Sui, Murat Yaman, Stefan Oehlers, Jordan A. Shavit, Vinitha A. Jacob

## Abstract

Endotoxemia is a feature of sepsis pathogenesis and has also been found to mediate the pathophysiology of multiple inflammatory conditions. In this work, we use a lipopolysaccharide (LPS) induced endotoxemia model in zebrafish to identify novel mediators of LPS toxicity. We performed transcriptomic studies on LPS-treated larvae, followed by *in silico* analysis, which revealed associations between the signatures of LPS-treated embryos and those of drugs involving diverse pathways. In parallel, we performed an *in vivo* screen using >1,500 FDA-approved compounds and identified multiple novel small molecules that reduced inflammation and prevented LPS toxicity. We focused on the direct thrombin inhibitor dabigatran, which was identified through both the *in vivo* and *in silico* analyses. We found that dabigatran co-administration significantly reduced the expression of inflammatory cytokines and completely protected zebrafish from endotoxemic death due to LPS. Surprisingly, we found that this protection occurs in prothrombin mutant fish, proving that protection from endotoxemia occurs independently of the anticoagulant function of dabigatran. We additionally found that dabigatran administration significantly decreased nitric oxide production and apoptosis compared to LPS treatment alone, suggesting possible mechanisms by which protection from endotoxemia is achieved. In summary, we identify several novel small molecules that prevent LPS-induced endotoxemia and show that one such small molecule, dabigatran, exerts a thrombin-independent effect on nitric oxide production and apoptosis. This and the other identified small molecules warrant further exploration in inflammatory conditions including sepsis.

## Introduction

Lipopolysaccharide (LPS), a cell wall component of gram-negative bacteria, induces an immune response upon recognition by Toll-Like Receptors (TLRs) ^1,2^. LPS accounts for the relative impermeability of the outer membrane of gram-negative bacteria to hydrophobic molecules, making them resistant to many antimicrobials ^3,4^. LPS-mediated endotoxemia mimics many features of sepsis pathogenesis and might also play a causative role in the pathophysiology of multiple chronic inflammatory disorders, including Parkinsons, Alzheimers, obesity, and atherosclerosis ^5^. Previous work in mammals suggests that the translocation of gut-derived LPS into the circulation might mediate some of these disease states ^6,7^. Additionally, LPS contributes to disease pathogenesis through its action as a prothrombotic molecule; LPS-induced inflammatory mediators activate procoagulant pathways, induce platelet activation, downregulate anticoagulant pathways and promote disseminated intravascular coagulation ^8,9^. Moreover, macrophages undergoing pyroptosis release tissue factor, further activating the coagulation cascade ^5,9^.

Given its role in multiple diseases, efforts have been made to block LPS biogenesis, neutralize endotoxin, and hinder LPS-mediated immune activation ^10–13^, with many of these attempts made in the context of sepsis. However, there is the possibility that therapeutics targeting LPS signaling can also mitigate the development of chronic inflammatory conditions such as those listed above.

In this work, we undertook large-scale *in silico* and *in vivo* analyses to identify novel small molecule mediators of LPS toxicity. Murine and other mammalian models are cost-prohibitive for the same types of large-scale *in vivo* screening studies. Zebrafish, on the other hand, allow for a speed of discovery not possible with mammals as they combine the scalability of *in vitro* systems with the complexity of a whole vertebrate organism.

In zebrafish, LPS-induced inflammation has been provoked either by injection into the embryonic yolk sac or immersion of embryos and larvae into LPS-containing media ^14–16^. Incubation with LPS-containing media allows uniform, rapid treatment of large numbers of fish. A response remarkably similar to that in mammals is observed, including inflammation, platelet activation, vascular leakage, and organ damage ^16^. Thus, many of the cardinal features of sepsis pathogenesis are recapitulated by this model.

We performed novel high throughput studies in a zebrafish model of LPS-provoked endotoxemia with the goal of identifying drugs that could be repurposed to combat acute and chronic inflammatory conditions. We conducted an *in silico* analysis, where we compared RNA-seq data from LPS-treated fish to publicly available RNA-seq datasets, revealing associations between LPS and drugs involving disparate pathways. In parallel, an *in vivo* pathway analysis was performed using >1,500 FDA-approved compounds to find those that mitigated LPS-induced endotoxemia, uncovering multiple novel small molecules. We focused on the sole anticoagulant (dabigatran) identified in both the *in silico* and *in vivo* analyses. Interestingly, the administration of dabigatran reduced the upregulation of inflammatory cytokines, significantly reduced nitric oxide production and apoptosis and abolished larval mortality from endotoxemia due to LPS. Surprisingly, this protection was found to occur independent of its inhibitory effect on thrombin. These data implicate dabigatran and other novel small molecules as anti-inflammatory agents that can be explored as potential therapeutics in acute and chronic conditions.

## Materials and Methods

### Animal care and husbandry

Zebrafish (*Danio rerio*) were raised in accordance with animal care guidelines as approved by the University of Michigan Animal Care and Use Committee. Wild-type fish were a hybrid line generated by crossing AB and TL strain zebrafish acquired from the Zebrafish International Resource Center. Genetic knockouts of prothrombin (*f2*) have been previously described ^17^. Tricaine (Western Chemical) was used for anesthesia and rapid chilling in an ice water bath for euthanasia.

### Small molecule treatments

Small molecule screens (MedChemExpress FDA drug repurposing set) were started at 3 dpf, where 3 larvae were plated per well in 96-well plates containing embryo media. All drugs were initially tested at a concentration of 25 μM for resistance to LPS toxicity, and confirmatory studies were performed with a larger group of fish (9-45) representing at least three different clutches (3 separate experiments), where the drugs were tested at various concentrations.

### Heart rate measurements

Heart rate was measured 3 hours after lethal LPS (40 μg/mL) administration and compared to untreated controls. Larvae were mounted in 0.8% low melting point agarose and visualized using an Olympus IX73 microscope. Heart rate was manually counted over 60 seconds for each larva. Each experiment consisted of at least 15 larvae. At least three different clutches (3 different experiments) were utilized.

### Real-time quantitative polymerase chain reaction (RT-qPCR)

RT-qPCR was performed as described previously ^18^. Total RNA was extracted from 3 dpf whole larvae 3 hours post treatment using the RNeasy Mini Plus kit (Qiagen) and transcribed into cDNA using Superscript III cDNA synthesis kit (Invitrogen). Transcript levels of genes of interest were quantified by qPCR (Mastercycler realplex^2^, Eppendorf) using Fast SYBR Green Mix (Applied Biosystems, Waltham, MA). Three equal pools of whole larvae (15-45 embryos per pool) were tested. Primer sequences are provided in Table S3. All expression data were normalized to the *actb2* gene, and significance was analyzed using the double delta C_t_ method as described ^19^.

### RNA-seq preparation and analysis

RNA-seq was performed as described previously ^18^. Three biological replicate pools of 30-45 3 dpf whole embryos were used per condition. Total RNA was extracted, and cDNA was prepared as above. Samples were sent to Novogene (Sacramento, CA) for further processing. Paired-end 150 base pair (bp) sequencing was obtained with the Illumina NovaSeq 6000 Sequencing System. Reads were mapped to exonic regions based on the zebrafish reference genome (GRCz11) using STAR36 (v.2.7.9a) and featureCounts37 (v.2.0.3). Differential expression analysis was conducted using the R package DESeq2 (^20^. All significant differentially expressed genes (FDR adjusted p-values) were converted to human genes using the R package babelgene. KEGG Pathway Analysis was then performed using the Enrichr platform, a publicly available tool ^21,22^. Overlap score is the number of significantly changed genes over the total number of genes in the pathway. The combined score considers the z-score (a value’s relationship to the mean) in addition to the adjusted p-value; this score has been found to outperform other ranking metrics ^22^. The combined score is used to assess the significance of gene set enrichment findings, with higher values being more significant; values > 30 have been considered significant in literature ^23,24^.

### Laser-mediated vascular endothelial injury

Larvae were treated with LPS and drug starting at 3 dpf and collected after 3 hours for assessment of time to clot formation, performed as previously described ^17,23,25,26^. Briefly, 3 dpf larvae were mounted in 0.8% low melting point agarose and visualized using an Olympus IX73 microscope with an attached Micropoint focusing system and pulsed-dye laser (Andor). For venous laser injury, 99 laser pulses were administered to the ventral surface of the posterior cardinal vein, 5 somites caudal to the anal pore. For arterial laser injury, 99 laser pulses were administered 5 somites caudal to the anal pore on the dorsal surface of the dorsal aorta. Time to occlusion (formation of an occlusive thrombus) was measured, where 120 seconds was the maximum time each larva was observed. Three biological replicates were performed for each experiment.

### In silico analysis

Differential gene expression data were retrieved from the RNAseq results as previously described ^18^. The human orthologues of the respective genes were obtained from the Zebrafish Information Network database (https://zfin.org/downloads/archive/2023.04.26) and the CLUE software platform was utilized in performing Connectivity Map-based analyses (https://www.sciencedirect.com/science/article/pii/S0092867417313090 (access date April 26, 2023)).

Genes were ranked based on the adjusted p-values for the gene expression level changes, and the top 150 upregulated or down-regulated genes were tested for the gene connectivity evaluations across 64 human cell lines and disparate perturbation profiles. The connectivity data were processed by using cmapR library in R (v4.2.2) environment (Natoli T (2023). cmapR: Cmap Tools in R. R package version 1.12.0, https://github.com/cmap/cmapR). The data were further filtered down to 27 cell lines of which most were hematopoietic/hepatic/pulmonary/non-tumorigenic origins, or endothelial phenotypes (A375, A549, ASC, BJAB, CD34, HA1E, HCC515, HEK293, HELA, HEPG2, HPTEC, HUH7, HUVEC, JURKAT, K562, MCF10A, NCIH508, NCIH596, NOMO1, OCILY19, OCILY3, PHH, THP1, TMD8, U2OS, U937, YAPC).

### Acridine Orange Staining

Larvae were incubated in 10 μg/ml Acridine Orange stain (Sigma) for 15 minutes in embryo media with LPS treatment and then rinsed three times before mounting in 0.8% agarose and imaged with an Olympus IX73 microscope. Quantification of intensity was performed in ImageJ.

### TUNEL Staining

The TUNEL assay was performed according to the instructions provided in the In Situ Cell Death Detection Kit, TMR red (Roche) for cryopreserved samples, with embryos cryopreserved in Optimal Cutting Temperature Compound (OCT) and cryosectioned prior to labeling. Images were quantified by an observer blinded to condition using a visual scoring system. The visual scoring system consisted of grades 0 through 3 (0 = 0-9% fluorescence in visual field, 1 = 10-20% fluorescence, 2 = 30-70% fluorescence, 3 = 80-100% fluorescence).

### NO Staining

Larvae were split into four treatment groups: control, LPS, dabigatran, and LPS with dabigatran. Each treatment group was simultaneously incubated with 5 uM of DAF-FM-DA solution in the dark at 25°C as previously described (Lepiller et al., 2007). After 2 hours larvae were washed with embryo media and mounted in 0.8% agarose. Imaging was done using an Olympus IX73 microscope. Fluorescence quantification was obtained utilizing ImageJ.

### Tail transection

Tg(lyzC:GFP)^nz117^ and TgBAC(mpeg1.1:EGFP)^vcc7^ transgenic larvae were anesthetized at 5 days post fertilization and tail transection was performed with a sterile scalpel blade. Wounded larvae were transferred into fresh embryo media with 25 μM dabigatran and incubated at 28.5 °C for 5 hours prior to imaging using a Nikon SMZ25 stereoscope. Leukocyte recruitment was quantified by measuring the fluorescent pixel area within 100 μm of the wound site in FIJI/ImageJ.

### Statistics

Statistics were performed using the GraphPad PRISM package. P<0.05 was considered significant. For qPCR and RNA-seq data, Student’s t-test on ΔΔC_t_ values was used to determine significance. For time to occlusion (TTO) data, Mann-Whitney *U* testing was used to determine significance.

### Study Approval

Approval for zebrafish studies was granted by the University of Michigan Institutional Animal Care and Use Committee, identification number PRO00010679 and the A*STAR Institutional Animal Care and Use Committee, identification numbers 211667 and 221694.

## Results

### LPS incubation results in inflammatory and coagulopathic changes in zebrafish embryos

*S. typhi* LPS was administered to zebrafish by incubation starting at 3 days post fertilization (dpf) in accordance with published data ^16^. A dose-dependent effect on lethality was noted over 24 hours. Hereafter, the lethal dose of LPS refers to the smallest amount that caused uniform lethality by 4 dpf, which was slightly different (ranging from 40-50 μg/mL) depending on the batch. A new dose-response curve was made for each batch of LPS tested, and an example is shown in Fig. 1A. In addition to increased mortality, heart rate was significantly decreased (Fig. 1B), and *grk5,* a gene involved in heart contractility, was significantly downregulated 3 hours after LPS administration (Fig. 1C).

**Figure 1:**
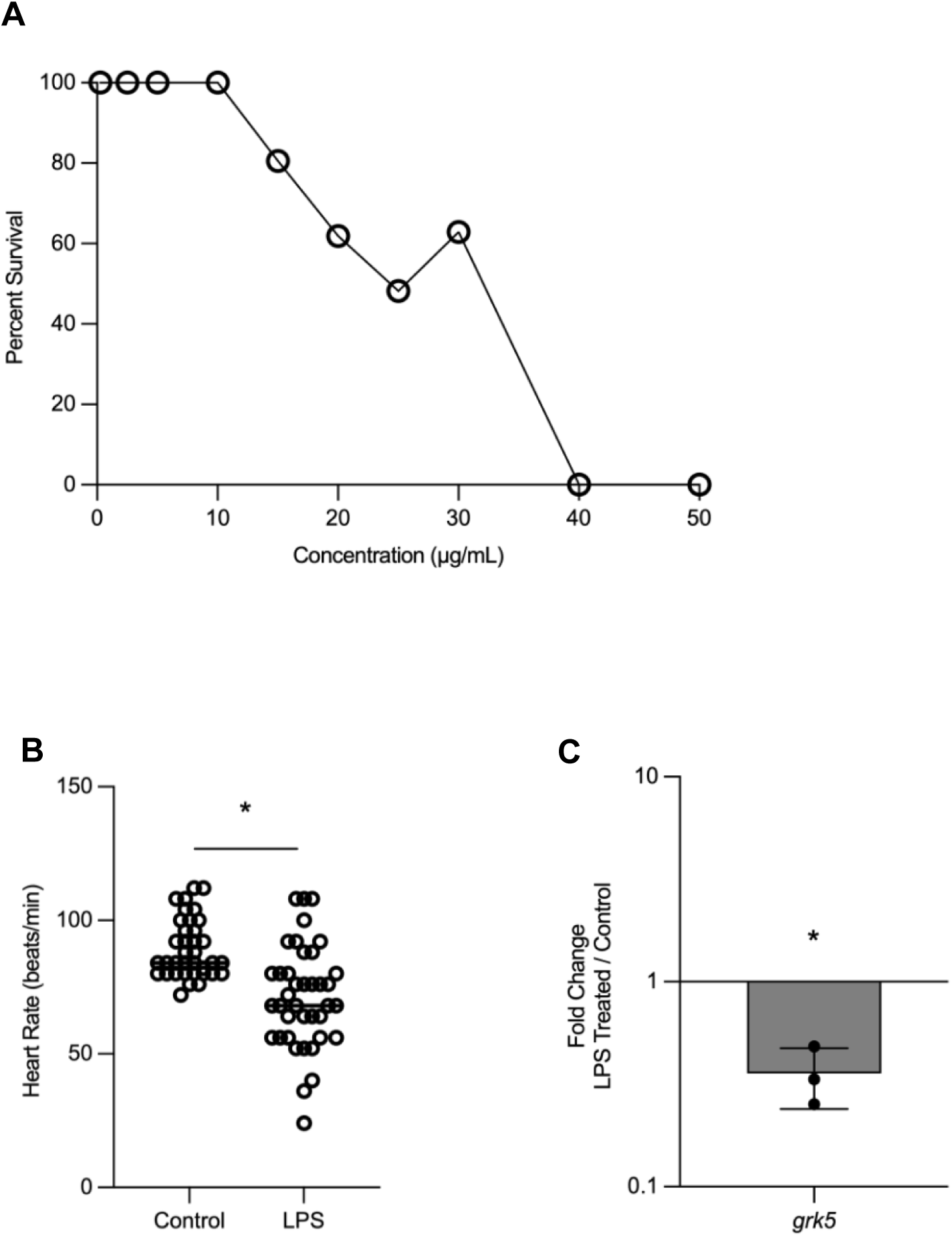
LPS incubation results in dose-dependent lethality and signs of end-organ damage in zebrafish larvae. (A) Percent survival of 4 days post fertilization (dpf) zebrafish larvae incubated with *S. typhi* LPS at 3 dpf. 3 clutches of 15 fish (total of 45) were used per concentration. (B) Heart rate (beats per minute) was counted 3 hours after LPS (40 μg/mL) administration and compared to controls. Each point represents a different fish, and the results include 3 separate experiments. (C) *grk5*, a gene involved in heart rate contractility, was measured by qPCR in LPS-treated fish compared to controls. Larvae from two or more clutches were pooled, where each point represents a separate clutch, and the average fold change in LPS-treated fish vs. controls was graphed. 30-45 total fish were used per time point. Bars indicate mean and standard deviation. Student’s t-test (two-tailed) on ΔΔC_t_ values was used to determine significance. *P<0.05.

To determine the significant biological pathways altered upon LPS administration, RNA-seq was performed on treated fish compared to controls as previously described (Guirgis et al., 2023). Kyoto Encyclopedia of Genes and Genomes (KEGG) pathway analysis was performed and revealed that “Pathogenic *E. coli* Infection” was the most significantly altered pathway in LPS-treated fish (Fig. 2A, complete analysis is shown in Table S1), confirming that administration induces a response akin to that provoked by bacteria. As expected, immune and interferon pathways such as TNF signaling were significantly changed (Fig. 2A). A subset of genes from these pathways were confirmed by qPCR and trended toward upregulation, with the genes *il1b*, *il6* and *irg1l* significantly upregulated (Fig. 2B). In addition to provoking a proinflammatory response, LPS is also known to be a prothrombotic molecule ^7,9^. RNA-seq data also showed that the complement, coagulation, and platelet activation pathways were significantly altered in LPS-treated zebrafish compared to controls (Fig. 2A). A subset of procoagulant genes from these pathways was upregulated, which was confirmed by qPCR (Fig. 2B). Interestingly, tissue factor b (gene *f3b*), but not tissue factor a (gene *f3a*) was significantly upregulated in LPS-treated fish compared to controls, suggesting possible arterial injury as *f3b* is thought to play a primary role in arterial, rather than venous, circulation ^26^. Time to occlusion (TTO) after venous and arterial laser-mediated endothelial injury was measured as previously described ^23,26^ and is illustrated in Fig. 2D. The TTO was found to be significantly increased in fish administered LPS for both systems (Fig. 2E, F). This change suggests a consumptive coagulopathy phenotype similar to what we have observed with the loss of natural anticoagulants ^23,27^.

**Figure 2:**
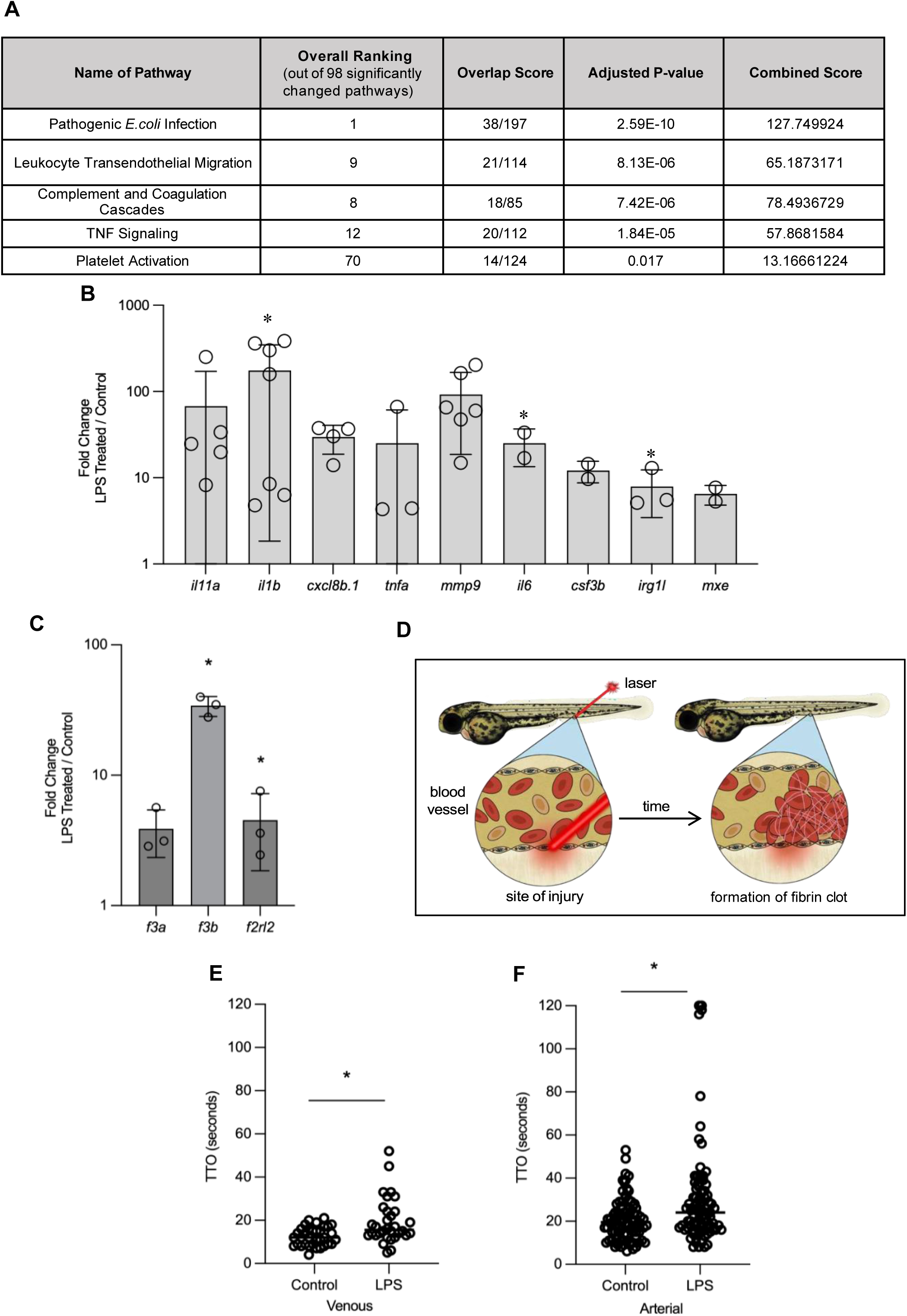
The immune and interferon signaling pathways and procoagulant pathways are significantly upregulated in LPS-treated fish compared to controls. (A) KEGG pathway analysis of RNA-seq data from whole LPS treated fish compared to controls. Significantly changed genes (adjusted p-value <0.05) from RNA-seq data on pooled 3 dpf LPS-treated vs. control fish (consisting of 30-45 embryos per category) performed in triplicate were used for further analyses. These genes were converted to human versions and then submitted for KEGG Pathway Analysis using the Enrichr platform, a publicly available tool developed by the Ma’ayan lab at Icahn School of Mount Sinai, NY (Chen et al., 2013, Kuleshov et al., 2016). Overlap score is the number of significantly changed genes over the total number of genes in the pathway. Combined score considers the z-score (a value’s relationship to the mean) in addition to the adjusted p-value. The combined score is used to assess the significance of gene set enrichment findings, with higher values being more significant. Adjusted p-value <0.05 is considered significant. Immune and interferon-related and complement and coagulation pathways are shown. For complete analysis, see Table S1. (B) A subset of genes in the immune and interferon pathways were tested by qPCR, using 3 dpf larvae treated with 40 μg/mL LPS vs. control conditions. Larvae from two or more clutches were pooled, where each point represents a separate clutch, and the average fold change in LPS-treated fish vs. controls is graphed. Bars represent mean and standard deviation. Student’s t-test (two-tailed) on ΔΔC_t_ values was used to determine significance. *P<0.05. (C) A subset of procoagulant genes was tested by qPCR, using 3 dpf larvae treated with 40 μg/mL LPS vs. control conditions. Larvae from two or more clutches were pooled, where each point represents a separate clutch, and the average fold change in LPS-treated fish vs. controls is graphed. Bars indicate mean and standard deviation. Student’s t-test (two-tailed) on ΔΔC_t_ values was used to determine significance. *P<0.05. A schematic of the laser injury assay used to measure time to occlusion is shown in (D), where the time to formation of an occlusive thrombus after laser pulses were delivered to the posterior cardinal vein or dorsal aorta was measured. 120 seconds was the maximum time each larva was observed. Time to occlusion (TTO) after laser injury in the venous (E) and arterial (F) circulation was measured in 3 dpf zebrafish larvae 3 hours after LPS administration and compared to controls. Each point represents a separate fish, and the observer was blinded to the treatment condition. P<0.05 is considered significant by Mann-Whitney *U* testing.

### In silico and in vivo analysis reveals small molecule mediators of LPS-mediated endotoxemia

To identify possible mediators of LPS toxicity, we initially undertook an *in silico* analysis comparing the RNA-seq signature of LPS-treated embryos to those of genes and compounds from publicly available databases (Table S2). After obtaining the list of small molecules related to LPS based on the gene expression profiles (Table S2), we further looked into mechanisms of action for each of these unique molecules as provided by the Connectivity Map (CMap) database. Even though some compounds did not have an annotated MoA, the data showed an intriguing pattern among the ones with detailed annotations. Notably, neuroactive compounds such as dopamine and serotonin receptor antagonists, along with ion channel and kinase inhibitors, emerged as the most commonly shared classes amongst the LPS-associated compounds (Table 1).

**Table 1:**
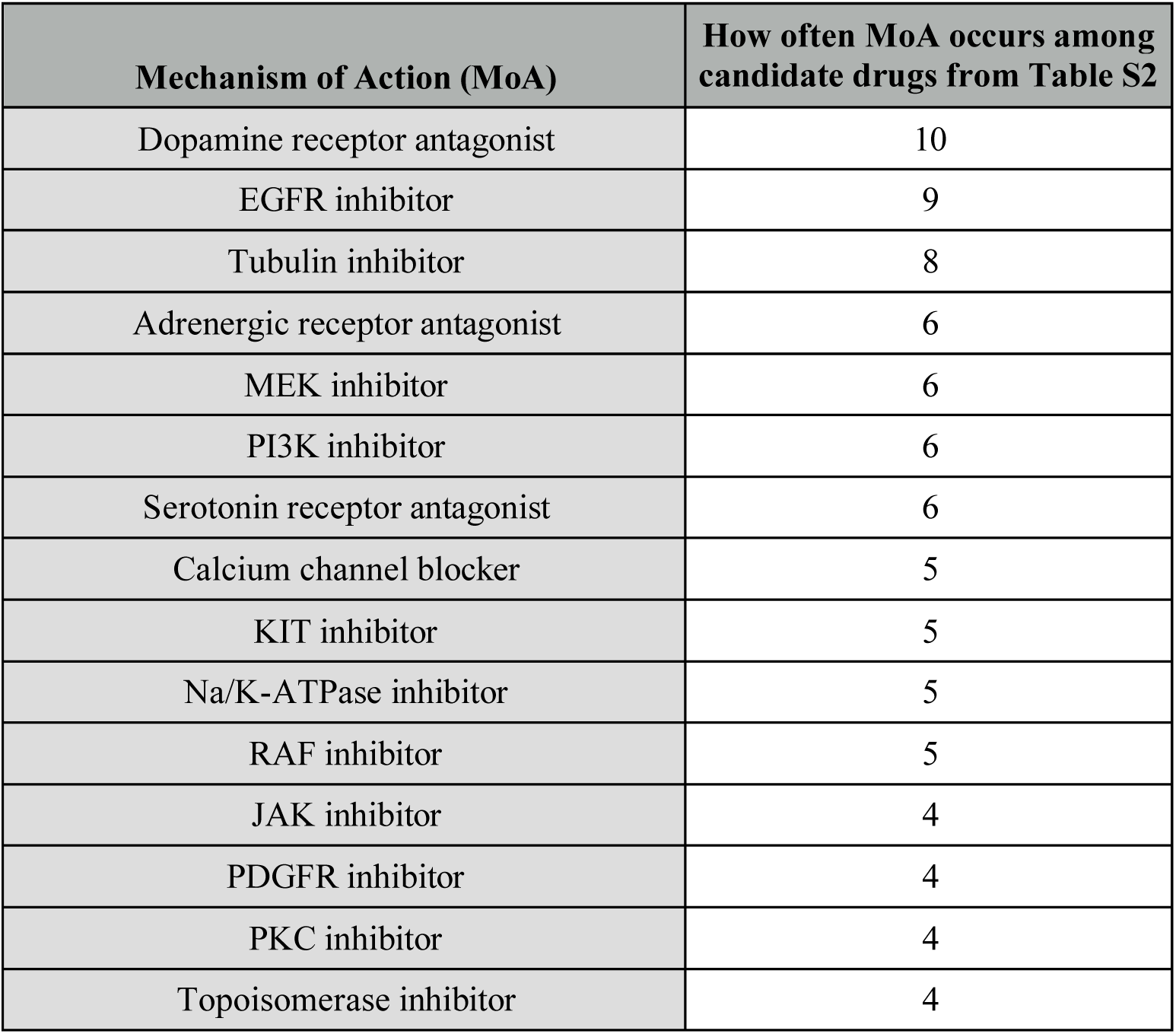
MoA of small molecules from Table S2 sorted by the number of times the MoA appears among the list of candidate drugs (only occurrences >= 4 are shown).

In parallel, we performed an *in vivo* analysis in zebrafish embryos. A lethal dose of LPS was co-administered at 3 dpf with an FDA-approved drug repurposing set consisting of >1500 compounds. Embryos were examined for survival 24 hours post-treatment, and each drug that was a ‘hit’ (i.e., enabled survival of at least one embryo) was validated in a larger cohort to confirm rescue. Approximately 20 small molecules were identified in this manner (Table 2). Most hits completely protected against LPS toxicity during the embryonic/early larval period, and others provided partial protection (Table 2, Fig. S1 A-T). These small molecule mediators of LPS toxicity spanned multiple categories, most heavily represented by serotonin and dopamine receptor modulators, chemotherapeutics, and anti-infectious agents. Similar to the *in silico* analysis, small molecules modulating the dopamine and serotonin pathways and those affecting tyrosine kinases were heavily represented *in vivo*. Four drugs were hits in both the *in silico* and *in vivo* analyses; these were amsacrine, conivaptan, vilazodone and dabigatran etexilate (the prodrug formulation of dabigatran, hereafter referred to simply as dabigatran) (Table S2, Table 2). Approximately half of the drugs identified *in vivo* (berberine, ketanserin, sunitinib, ceritinib, fedratinib, nelfinavir, saquinavir, indacaterol, dabigatran) have been shown to ameliorate LPS effects, and to function as mammalian anti-sepsis agents ^28–48^, suggesting conservation between zebrafish and mammals in terms of the effects of these small molecules. The remaining hits from the *in vivo* screen – ziprasidone, pergolide, vilazodone, bedaquiline, rucaparib, pralsetinib, amsacrine, conivaptan, deserpidine, and lomitapide – are, to our knowledge, novel mediators of LPS toxicity (Table 2). Despite some variability, there was an overall trend toward decreased inflammatory cytokine production when a subset of these drugs were co-administered with LPS and compared to LPS treatment alone (Fig. S2 A-L). Thus, these novel mediators of LPS toxicity may function as anti-inflammatory agents.

**Table 2:** Small molecules protecting zebrafish embryos from LPS toxicity.

| Drug Category | Drug identified from current zebrafish LPS screen | Main Mechanism of Action | Drug previously tested with LPS in mammals?/ Novel | Drug previously tested in sepsis?/ Novel |
| --- | --- | --- | --- | --- |
| **Dopamine and Serotonin Modulators** | Ziprasidone | Dopamine and serotonin antagonism | Novel | Novel |
|  | Pergolide | Dopamine receptor agonist | Novel | Novel |
|  | Vilazodone | Inhibits serotonin transporter, partial agonist at serotonin receptors | Novel | Novel |
|  | Brexipiprazole | Dopamine and serotonin receptor modulation | Novel | Novel |
|  | Ketanserin | Serotonin receptor antagonist | Mouse (Liu et al., 2013, Sugino et al., 2009) | Clinical trial for septic shock (Vellinga et al., 2015) |
| **Chemotherapeutics** | Rucaparib | Inhibits poly(ADP-ribose) polymerase | Novel | Novel |
|  | Sunitinib | Inhibits receptor tyrosine kinases | Mouse (Zhao et al., 2019) | Novel |
|  | Fedratinib | Inhibits janus receptor kinase 2 | Human macrophages (Akcora et al., 2019) | Novel |
|  | Pralsetinib | Rearranged during transfection kinase inhibitor | Novel | Novel |
|  | Ceritinib | Inhibits autophosphorylation of anaplastic lymphoma kinase | Human endothelial cells (Dayang et al., 2021, Hu et al., 2019) | CLP induced sepsis in mice (Hu et al., 2019) |
|  | Amsacrine | DNA intercalator | Novel | Novel |
| **Antiviral/ Antibacterial Agents** | Nelfinavir | Protease inhibitor | Human macrophages (Wallet et al., 2012) | CLP induced sepsis in mice (Weaver et al., 2004) |
|  | Saquinavir | Protease inhibitor | Mouse (Yao et al., 2023, Tong et al., 2021), Human lung cells (Zhang et al., 2018) | CLP induced sepsis in mice (Tong et al., 2021) |
|  | Ledipasivir | Inhibits viral phosphorylation | Novel | Novel |
|  | Bedaquiline | Blocks proton pump for ATP synthase | Novel | Novel |

**Table 2 continued: Small molecules protecting zebrafish embryos from LPS toxicity**
| Drug Category | Drug identified from current zebrafish LPS screen | Main Mechanism of Action | Drug previously tested with LPS in mammals?/ Novel | Drug previously tested in sepsis?/ Novel |
| --- | --- | --- | --- | --- |
| Miscellaneous | Indacaterol | β-adrenoreceptor agonist | Guinea pigs (Aritake et al., 2021), Human lung macrophages (Maarsingh et al., 2021) | Novel |
|  | Conivaptan | Vasopressin receptor antagonist | Novel | Novel |
|  | Deserpidine | Vesicular monoamine transporter blocker | Novel | Novel |
|  | Lomitapide | Microsomal triglyceride transfer protein inhibitor | Novel | Novel |
|  | Dabigatran | Thrombin inhibitor | Mouse (Kimura et al., 2020) | Mouse sepsis models, including CLP (Lei et al., 2021), *S. aureus* bacteremia in humans (Butt et al., 2021) |
|  | Berberine | Activates AMPK (increases glycolysis) | Mouse (Chen et al., 2022), rats (Chen et al., 2021, Zhao et al., 2023), mouse macrophage cells (Zhao et al., 2023), bovine endometrial epithelial cells (Fu et al., 2023) | CLP-induced sepsis in mice (Chen et al., 2023), cecal slurry-induced sepsis (Li et al., 2021), *S. aureus*-induced septic arthritis in mice (Asila et al., 2022) |

### Dabigatran reduces LPS mediated inflammation and mortality

Given the availability of tools to study coagulation in zebrafish ^25,49^ we decided to characterize further the one anticoagulant, dabigatran, that was identified through both *in silico* and *in vivo* analyses. Although a prior study has shown only a modest effect of dabigatran on lethality in sepsis models ^39^, we found that it completely averted LPS-induced lethality (Fig. 3A). Expression of inflammatory cytokines and procoagulant genes were downregulated in fish co-treated with LPS and dabigatran compared to LPS alone (Fig. 3B). As previously shown ^17^, time to venous occlusion was increased in dabigatran treated fish, indicating successful anticoagulation (Fig. 3C). There was no additive effect on occlusion with co-treatment.

**Figure 3:**
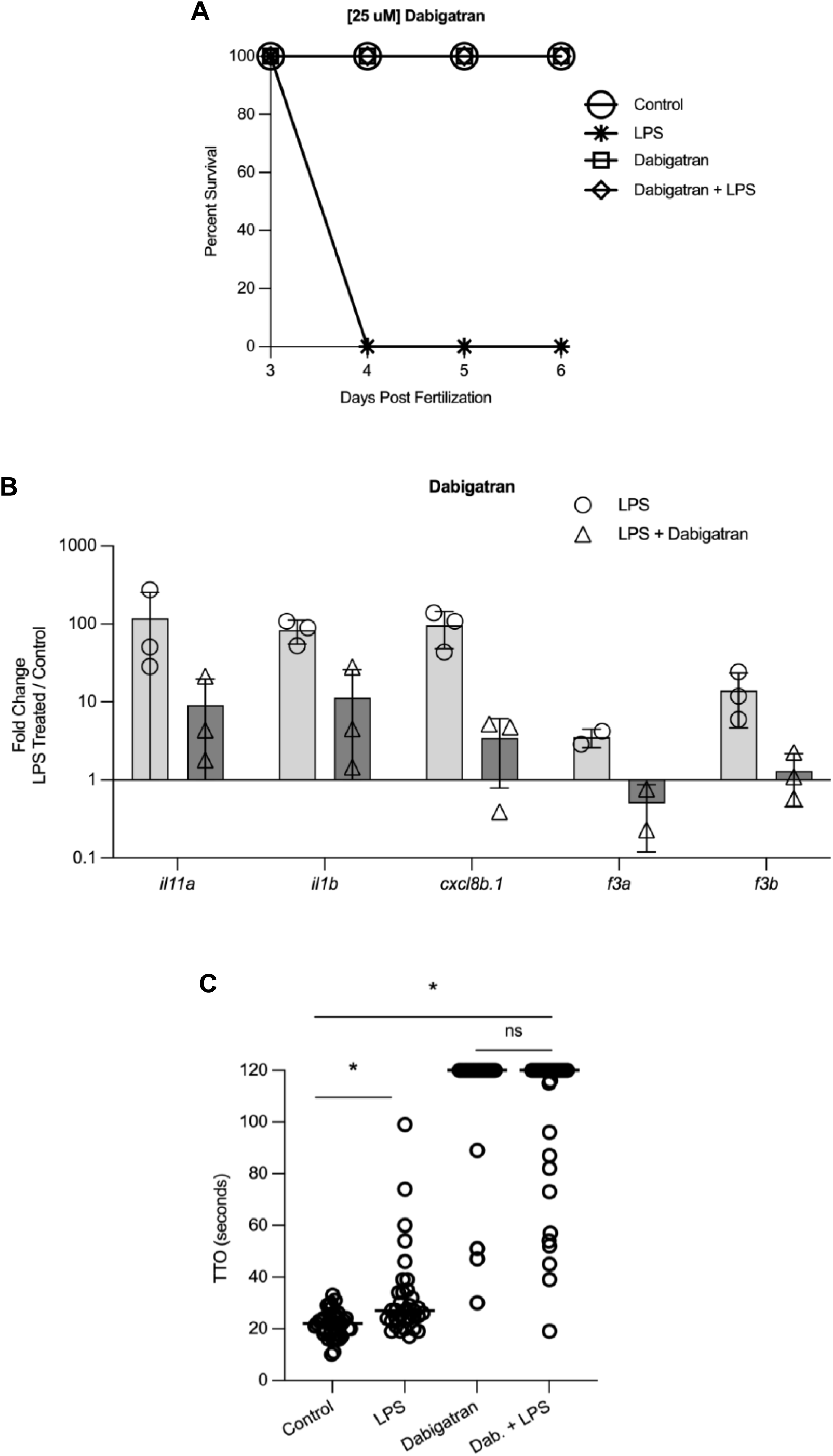
The anticoagulant dabigatran mitigates LPS-induced lethality and reduces the expression of inflammatory genes. (A) Zebrafish were administered a lethal dose of LPS with 25 μM dabigatran along with appropriate controls and survival was tracked daily from 3 through 6 dpf. (B) Expression of inflammatory genes was evaluated by qPCR 3 hours after drug administration in 3 dpf larvae treated with LPS and dabigatran. Pooled larvae (15-45) from a clutch were used, where each point represents a separate clutch, and the average fold change in LPS-treated fish vs. controls is graphed. Bars indicate mean and standard deviation. Student’s t-test (two-tailed) on ΔΔC_t_ values was used to determine significance. *P<0.05. (C) Time to occlusion (TTO) after laser injury in the venous circulation was measured in 3 dpf zebrafish larvae 3 hours after LPS and dabigatran administration. Each point represents a single larva, and the observer was blinded to the condition. P<0.05 was considered significant by Mann-Whitney *U* testing.

We next determined whether dabigatran administration could produce a more general anti-inflammatory effect. Tail transection, an established method to induce inflammation, was performed on 3 dpf larvae. We found that dabigatran did not reduce neutrophil or macrophage recruitment to the tail transection site, suggesting that its protection might be more specific to LPS rather than representing a general anti-inflammatory response (Fig. S3).

### Dabigatran protects zebrafish from LPS toxicity independent of its anticoagulant function

An anti-sepsis effect of dabigatran mediated through fibrinogen-like protein 2 (Fgl2) has been described^39^. To determine whether the protection from LPS toxicity was dependent on the anticoagulant effect of dabigatran, we evaluated three other commonly used anticoagulants – two affecting factor Xa, rivaroxaban and apixaban, as well as another thrombin inhibitor, argatroban. Rivaroxaban and apixaban were previously shown to be active in zebrafish and were used here at the same concentration ^17^. However, we required shorter incubation times due to the rapid lethality induced by LPS. As expected, all anticoagulants increased the TTO (Fig. 4A, Fig. S4 A, B). Interestingly, some of the anticoagulants appeared to have a synergistic effect with LPS, significantly increasing the time to occlusion when compared to the drug alone (Fig. 4A, Fig. S4 A, B). However, none of these conferred protection from LPS toxicity (Fig. 4B, Fig. S4 C, D), suggesting that anticoagulation might not be the mechanism through which dabigatran mitigates LPS-induced endotoxemia.

**Figure 4:**
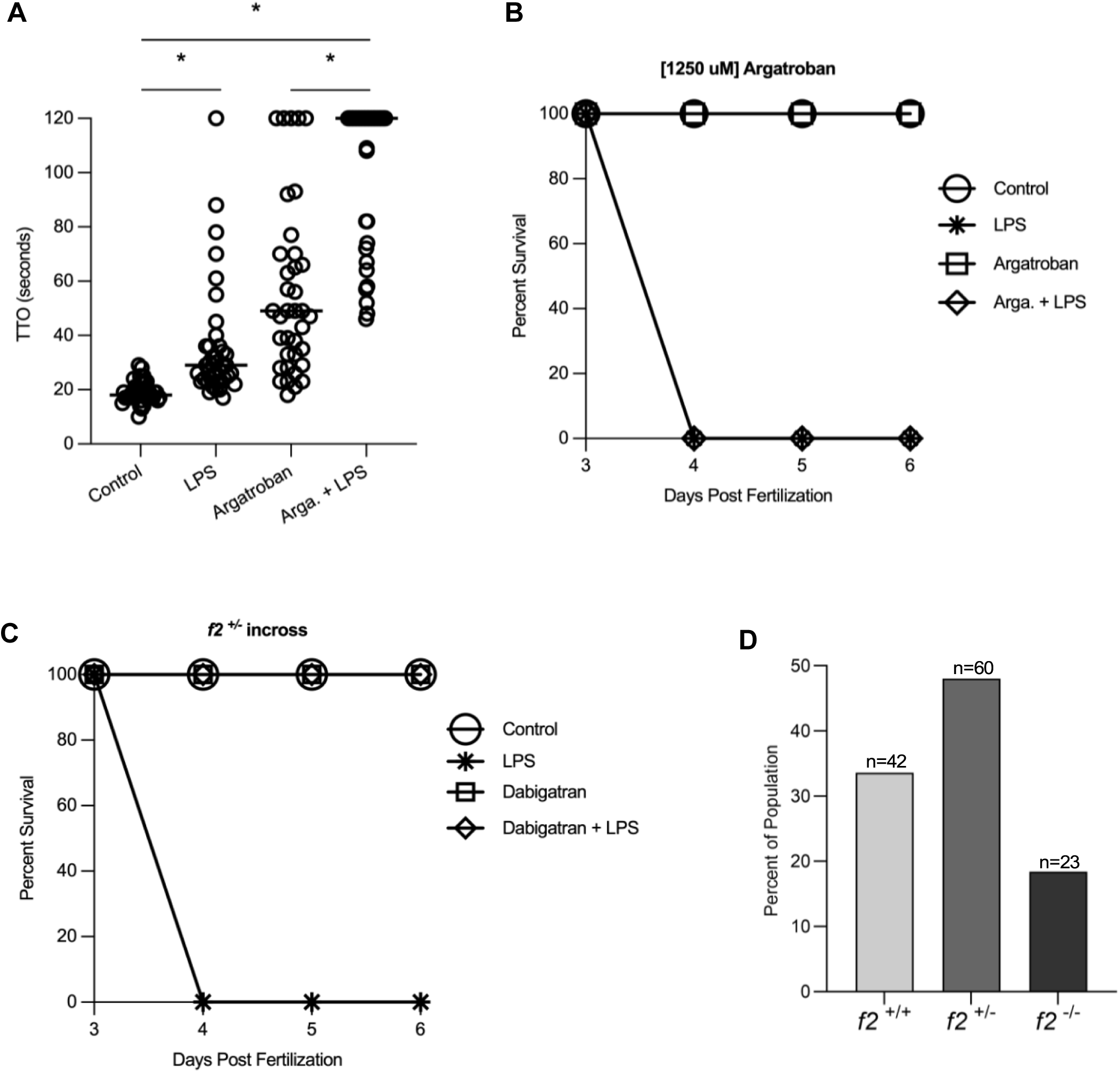
Dabigatran protection from LPS toxicity occurs independently of prothrombin. (A) Time to occlusion (TTO) after laser injury in the venous circulation was measured in 3 dpf zebrafish larvae 3 hours after LPS and argatroban administration. Each point represents a single larva, and the observer was blinded to the condition. P<0.05 was considered significant by Mann-Whitney *U* testing. (B) Zebrafish were treated with argatroban and/or LPS at 3 dpf and observed daily until 6 dpf for survival. (C) Three dpf larvae from *f2* heterozygous incrosses were divided into four groups: untreated controls, 40 μg/mL LPS, 25 μM dabigatran, and LPS/dabigatran. All groups were followed for 3 days for survival. (D) Genotype distribution of surviving larvae.

To further test this hypothesis, we employed a prothrombin knockout background (null mutation in the *f2* gene ^17^. Larvae from *f2* heterozygous incrosses were divided into four groups: untreated controls, LPS, dabigatran, and LPS + dabigatran. Each group was followed for survival over 3 days. LPS alone was lethal in all genotypes (Fig. 4C), revealing that thrombin inhibition is not necessary for protection from LPS. The presence of homozygous mutants was confirmed by genotyping all clutches, except for LPS alone due to their rapid degradation (Fig. 4D). Thus, dabigatran administration protected *f2* homozygous mutants from lethal LPS, proving that its mechanism of protection occurs independently of thrombin inhibition.

### Dabigatran administration reduces LPS mediated nitric oxide production and apoptosis

Nitric oxide (NO) production can contribute to the hypotension seen in septic shock ^50^, and LPS has previously been showed to increase NO ^50^. We confirmed that LPS administration increased NO production in our system (Fig. 5A, B). Additionally, dabigatran has previously been found to suppress nitric oxide synthase ^51^. We therefore tested whether dabigatran treatment could reduce NO production and found this to be the case (Fig. 5A, B).

**Figure 5:**
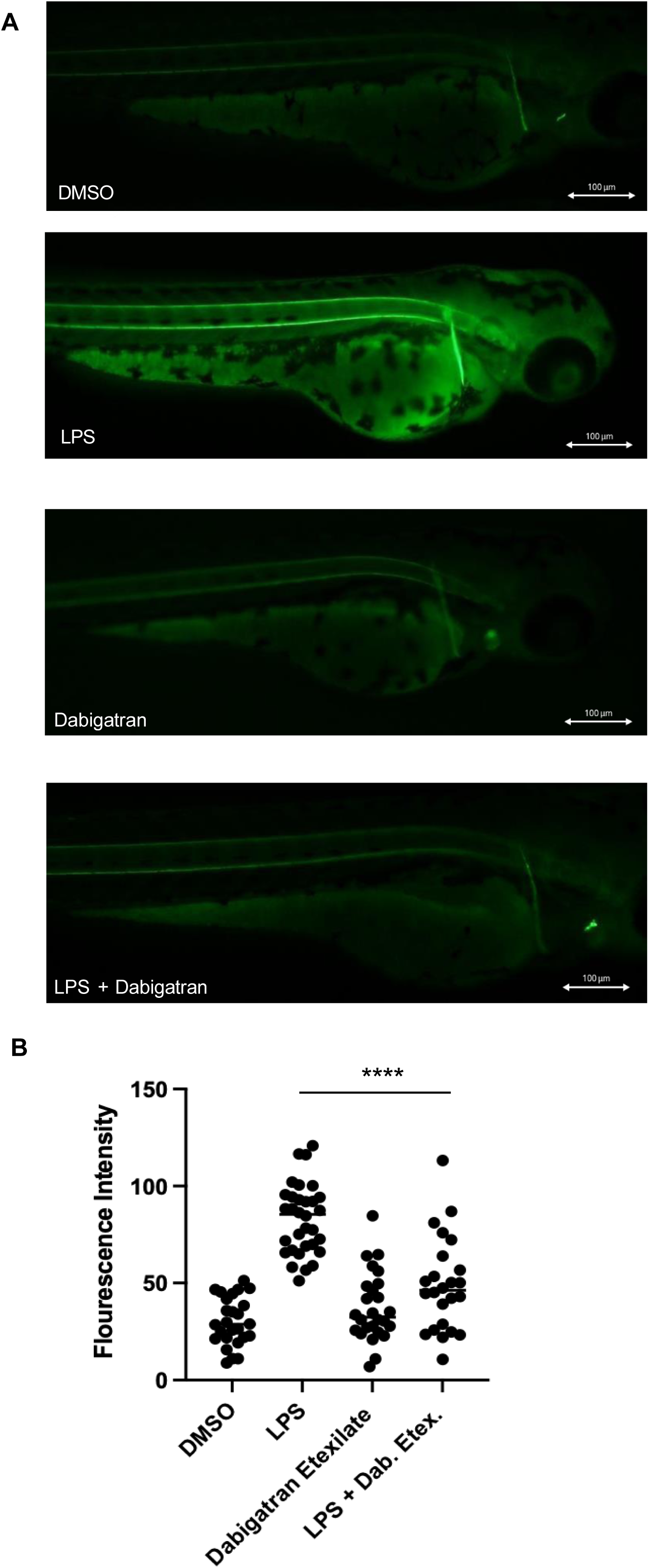
Dabigatran reduces LPS mediated nitric oxide production. (A) Representative images of 3 dpf fish stained with DAF-FM DA to visualize nitric oxide production after the indicated treatment. (B) ImageJ analysis showing mean fluorescence intensity of images of 3 dpf DAF-FM DA stained fish with the indicated treatment; observers were blinded to condition. Each point represents a different fish, and the results include 3 separate experiments. Student’s t-test (two-tailed) on ΔΔC_t_ values was used to determine significance. *P<0.05.

RNA sequencing data suggested that apoptosis related pathways were increased in LPS-treated fish (Fig. 6A). As NO production has been shown to lead to apoptosis after LPS exposure ^52^, we next tested whether this represented a mechanism by which protection from LPS is conferred. We found that cell death (Fig. 6B, C) and apoptosis (Fig. 6D, E) were significantly decreased in dabigatran and LPS co-treated fish. These data suggest that decreased NO production and apoptosis might be mechanisms by which dabigatran protects against LPS toxicity.

**Figure 6:**
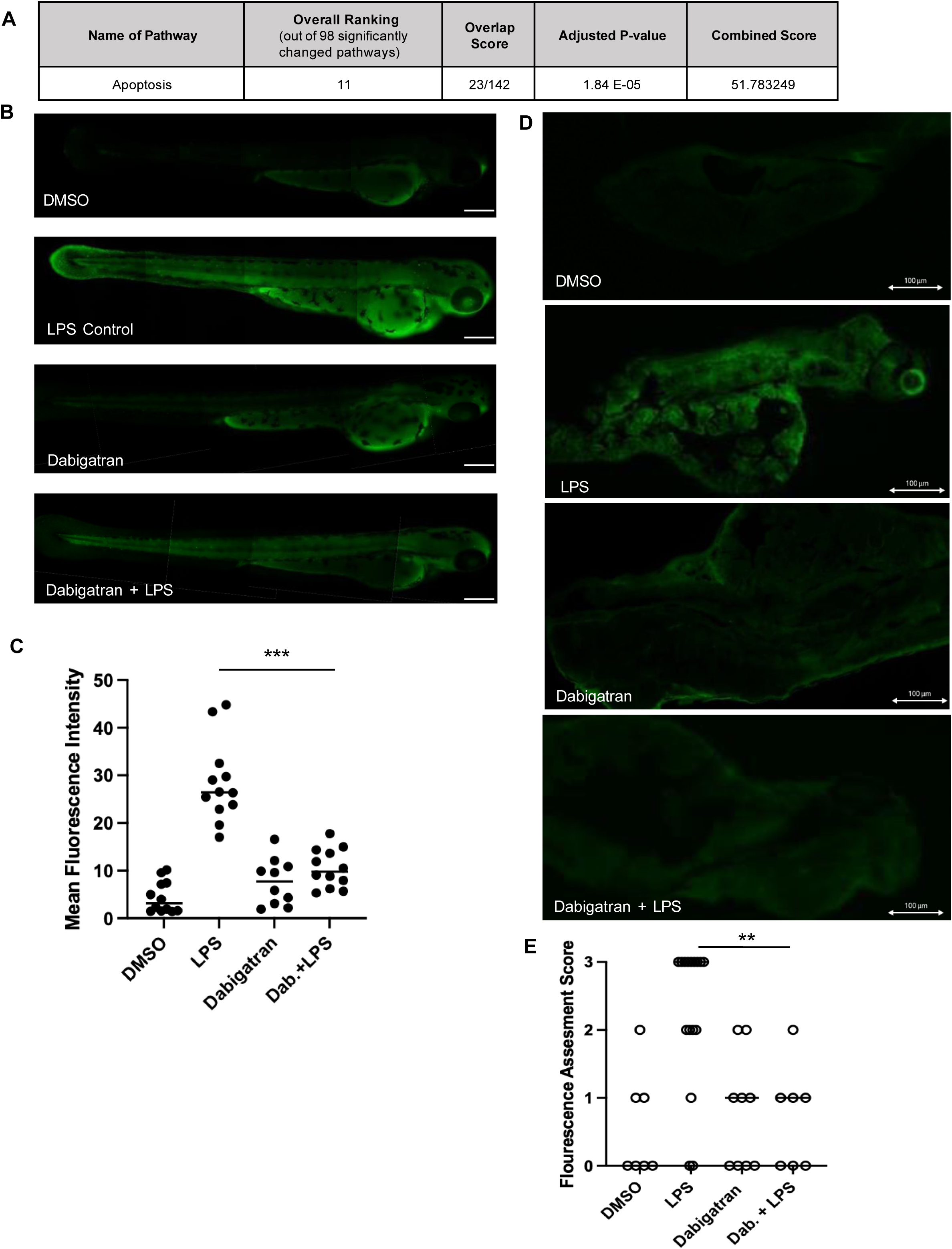
Dabigatran reduces LPS mediated cell death and apoptosis. (A) KEGG pathway analysis of RNA-seq data from whole LPS treated fish compared to controls. Significantly changed genes (adjusted p-value <0.05) from RNA-seq data on pooled 3 dpf LPS-treated vs. control fish (consisting of 30-45 embryos per category) performed in triplicate were used for further analyses. These genes were converted to human versions and then submitted for KEGG Pathway Analysis using the Enrichr platform, a publicly available tool developed by the Ma’ayan lab at Icahn School of Mount Sinai, NY (Chen et al., 2013, Kuleshov et al., 2016). Overlap score is the number of significantly changed genes over the total number of genes in the pathway. Combined score considers the z-score (a value’s relationship to the mean) in addition to the adjusted p-value. The combined score is used to assess the significance of gene set enrichment findings, with higher values being more significant. Adjusted p-value <0.05 is considered significant. Apoptosis related pathways are shown. For complete analysis, see Table S1. (B) Representative images of 3 dpf fish stained with acridine orange after the indicated treatment. (C) ImageJ analysis showing mean fluorescence intensity of images of 3 dpf acridine orange stained fish with the indicated treatment; observers were blinded to condition. Each point represents a different fish, and the results include 3 separate experiments. Student’s t-test (two-tailed) on ΔΔC_t_ values was used to determine significance. *P<0.05. (D) Representative images of 3 dpf fish stained with TUNEL after the indicated treatment. (E) Apoptosis was scored in 3 dpf TUNEL stained fish with the indicated treatement; observers were blinded to condition. Each point represents a different fish, and the results include at least 2 separate experiments. Student’s t-test (two-tailed) on ΔΔC_t_ values was used to determine significance. *P<0.05.

## Discussion

We have identified several small molecule mediators of LPS toxicity that can act as anti-inflammatory agents. Approximately half had been previously explored in mammalian LPS models, and all have been shown to reduce inflammation and ameliorate toxicity ^28–31,33–37,40,42,43,47,48,53–55^, suggesting strong concordance between zebrafish and mammals. These mediators were found to act through multiple mechanisms, including inhibition of TLR and NFκB activation ^33,54^ as well as inhibition of inflammation through NFκB-independent mechanisms ^34,35^. Other potential mechanisms include effects on apoptosis ^30^ and NOS (nitric oxide synthase) activation ^31^. A smaller number of the identified drugs have also been investigated in mammalian sepsis models ^32,39,41,43–45,47,56^, all with beneficial effects, indicating further concordance with our zebrafish data. The novel mediators of LPS toxicity that have not previously been tested in other studies – amsacrine, bedaquiline, conivaptan, deserpidine, lomitapide, pergolide, prasetinib, rucaparib, vilazodone, ziprasidone – have as yet undetermined mechanisms of action and represent avenues for further research in both mammalian inflammation and sepsis.

Clinical trials using anticoagulants in the context of sepsis have proven equivocal at best ^57–66^, although it is still possible that specific subtypes of sepsis patients may benefit from anticoagulant therapy. Previous studies have shown that dabigatran reduces inflammation (^39,40^ and can enhance survival in mouse models of sepsis ^39^, congruent with a cohort study revealing that patients on dabigatran had a lower incidence of *Staphylococcus aureus* bacteremia ^41^. Dabigatran has also been found to mitigate inflammation in multiple contexts, including diet ^67^, oxysterols ^68^, and acute myocardial infarction ^51^. However, it is unclear whether these beneficial effects were due to the anticoagulant function of dabigatran. Our data suggest that dabigatran reduces inflammation and promotes survival from LPS toxicity through a novel mechanism independent of its direct anticoagulant effect on thrombin.

We found that dabigatran administration reduces LPS mediated NO production and apoptosis, suggesting that this is one possible mechanism by which dabigatran confers protection from LPS. Dabigatran has previously been shown to suppress nitric oxide synthase in acute myocardial infarction ^51^ and future experiments will clarify the mechanism by which dabigatran reduces LPS mediated nitric oxide production.

There are several limitations of this study. Due to the stringency of the screen (mortality), it is likely that other small molecules conferring beneficial effects but not affecting mortality (e.g., those that reduce inflammation) were not identified. This study also does not necessarily identify the mechanism by which each drug acts to reduce LPS toxicity. It is possible that the effect is mediated through a non-primary or previously unknown mechanism of action (as is the case with dabigatran).

Our data suggest that the novel small molecule mediators of LPS toxicity uncovered through our screen can be explored as anti-inflammatory and anti-sepsis agents in mammals. Given that the hits previously examined in mammalian systems were also found to have beneficial effects in this zebrafish model, we expect that at least some of the novel drugs will translate to mammalian models.

Previous work and our data suggest that dabigatran can act as an anti-inflammatory and potential anti-sepsis agent ^39–41^. We show here definitive proof that dabigatran acts independently of thrombin to exert its rescue of lethal endotoxemia. Venous thromboembolism prophylaxis is the standard of care for most adult hospitalized inpatients, especially those admitted for sepsis ^69,70^. The most common agents used are unfractionated and low molecular weight heparins, but the data from our study suggest the possibility that dabigatran could be an alternative with a dual benefit –both anticoagulant and completely distinct anti-LPS functions. Randomized controlled trials in human inflammatory conditions or inpatients would be relatively easy to support given the successful use of direct oral anticoagulants and the indication for anticoagulation in hospitalized patients ^69,70^.

## Acknowledgments

A*STAR IMCB Aquarium Platform for expert zebrafish husbandry.

## Data Availability

RNAseq data have been submitted to NCBI Gene Expression Omnibus and are available under accession number GSE245299. Any other original data can be obtained by contacting the corresponding author at.

## Author Contributions

V.J., J.A.S., S.O. and A.F. conceived and planned experiments. V.J., A.F., R.M., W.W. and X.S. performed and analyzed all experiments and M.Y. performed *in silico* analysis. V.J. wrote the manuscript, with input from all authors. J.A.S. revised the manuscript.

