## Supplementary figures and images for "Dabigatran prevents lipopolysaccharide mediated apoptosis in zebrafish through a thrombin independent mechanism"

### Supplemental Figures

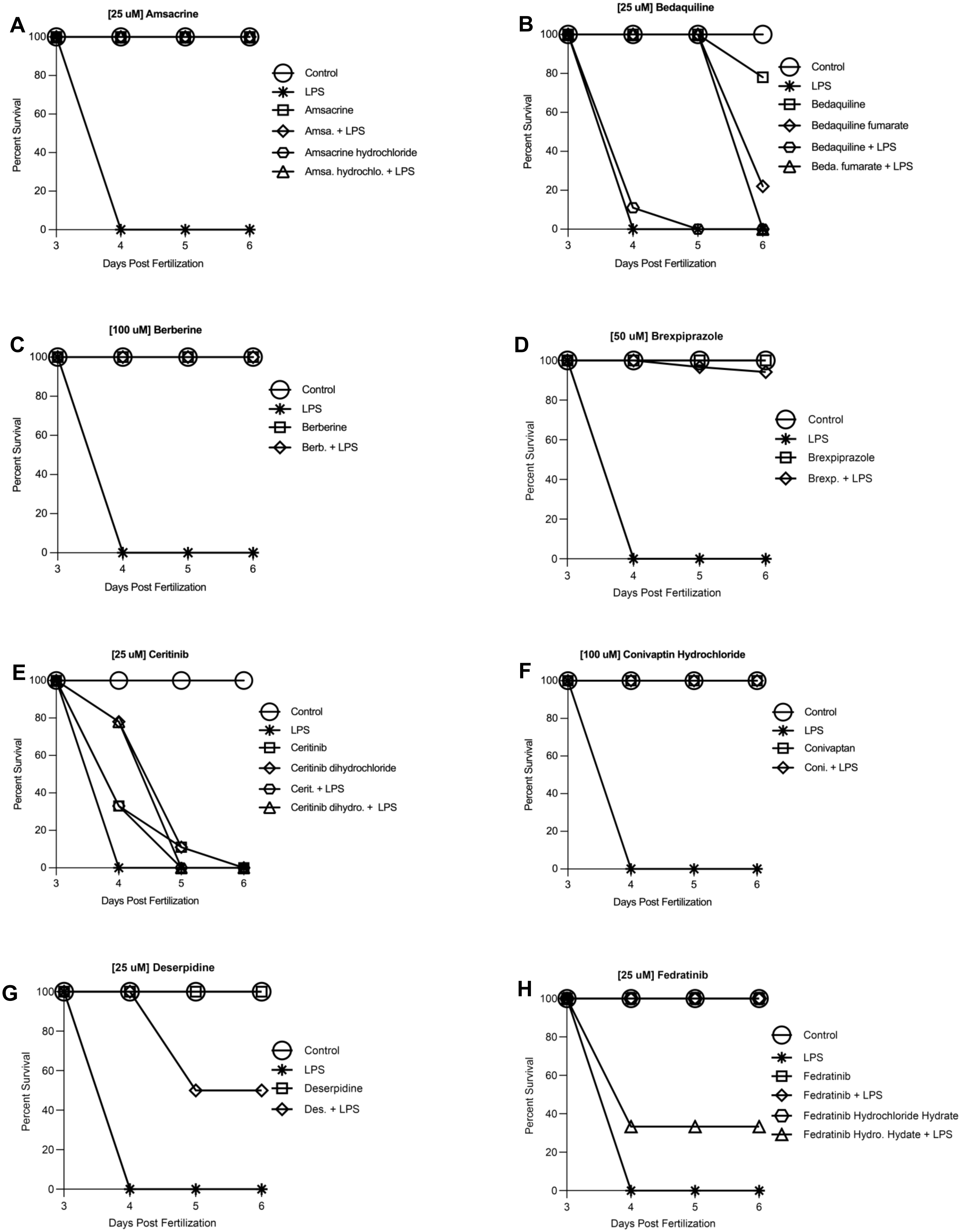

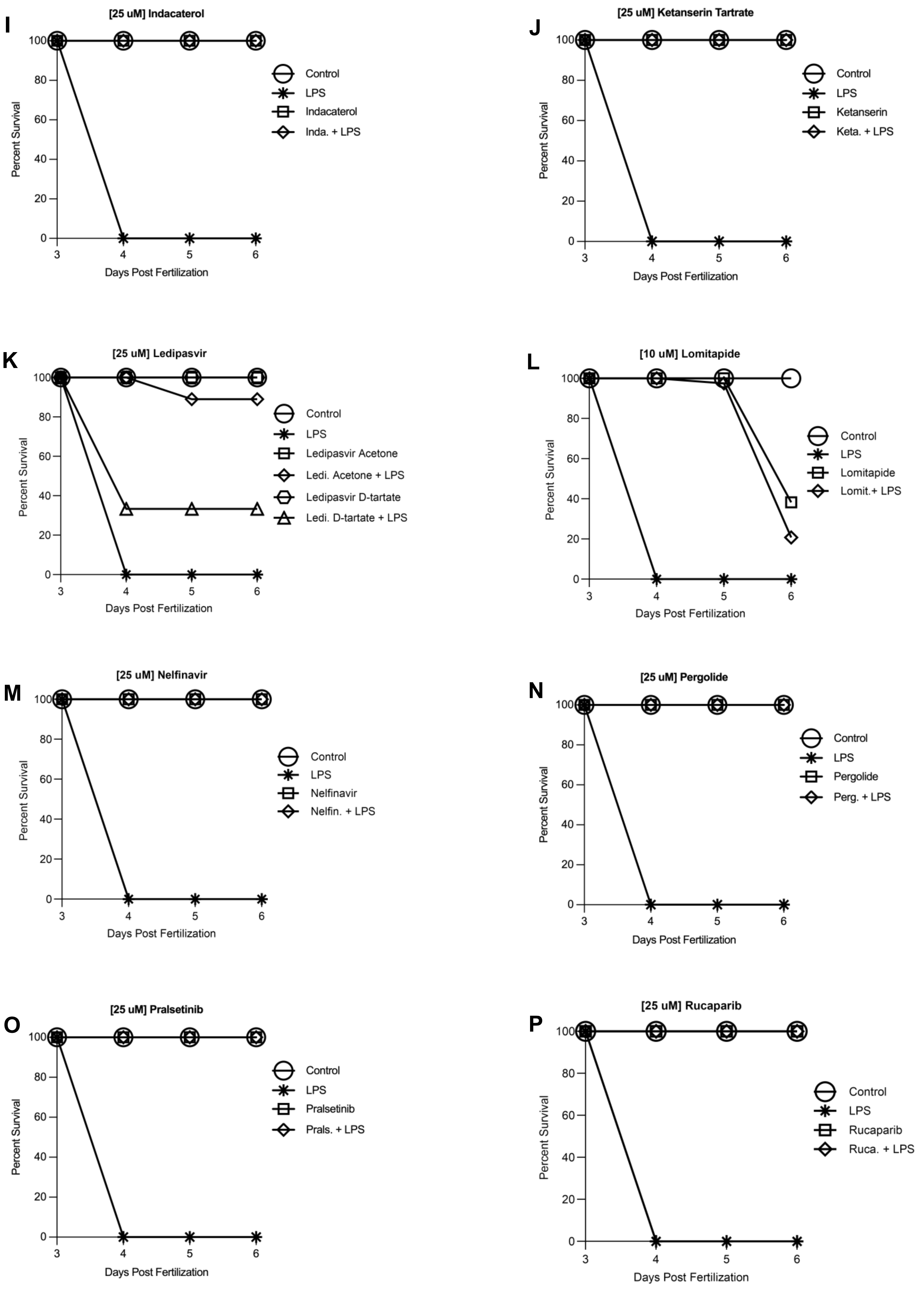

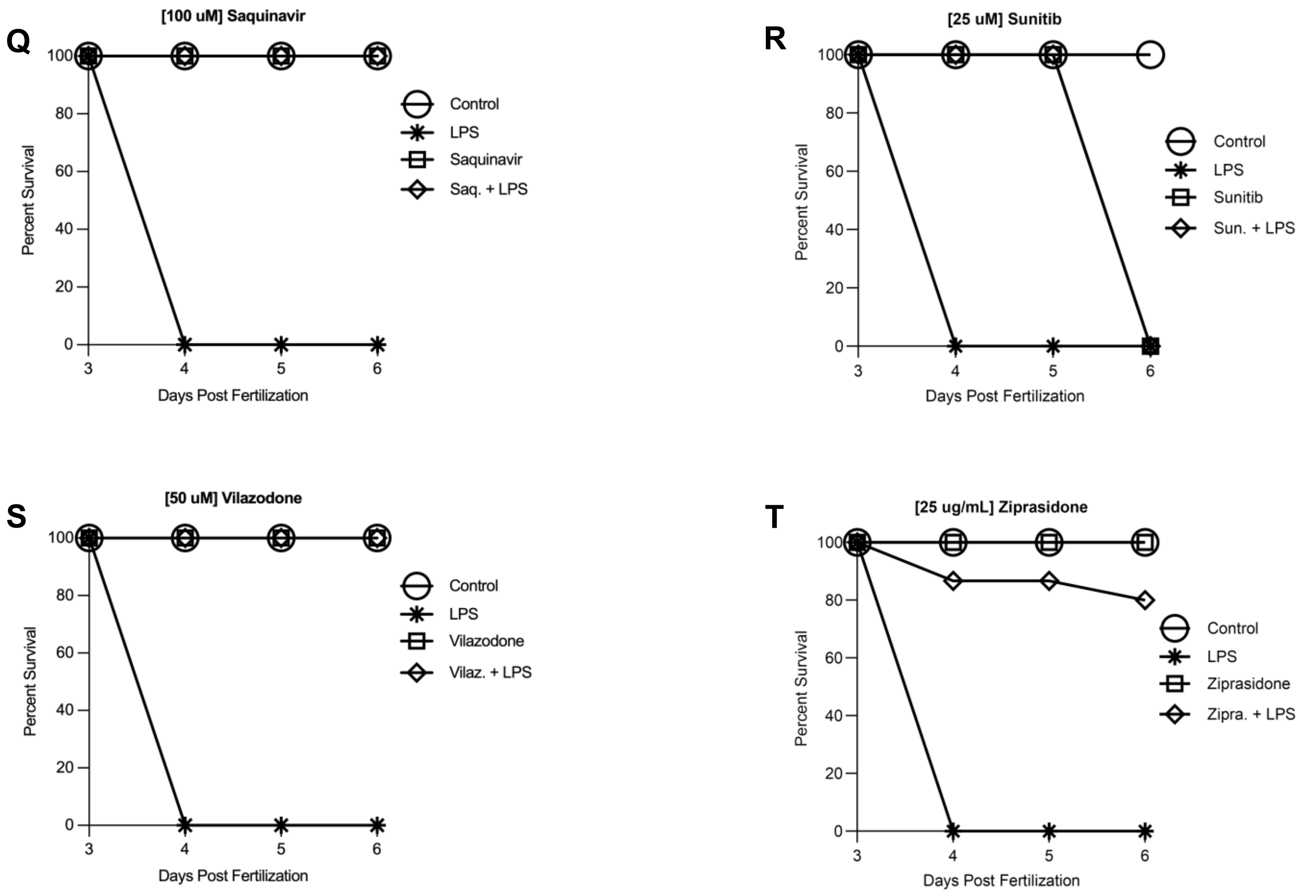

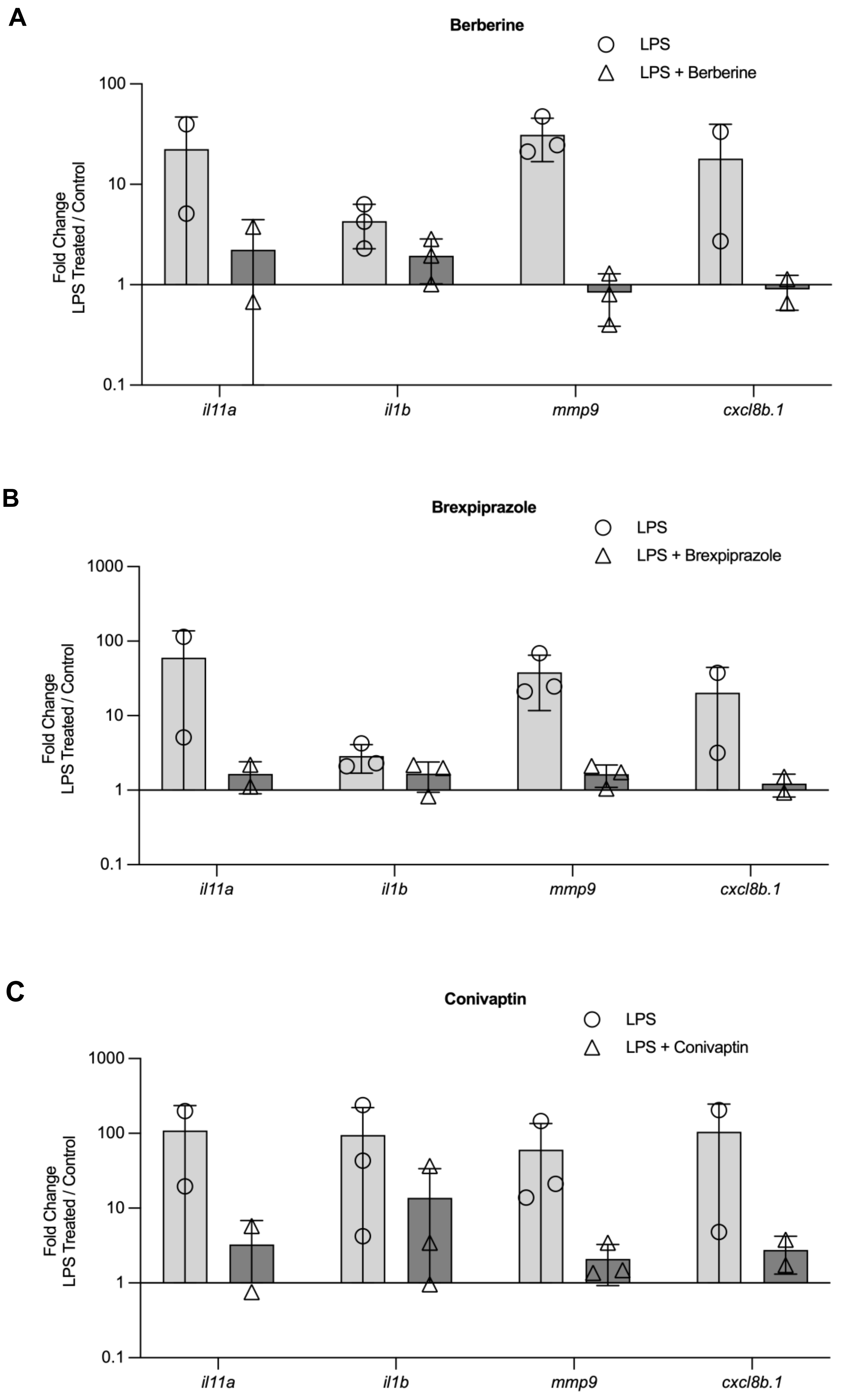

**Figure S2**

D

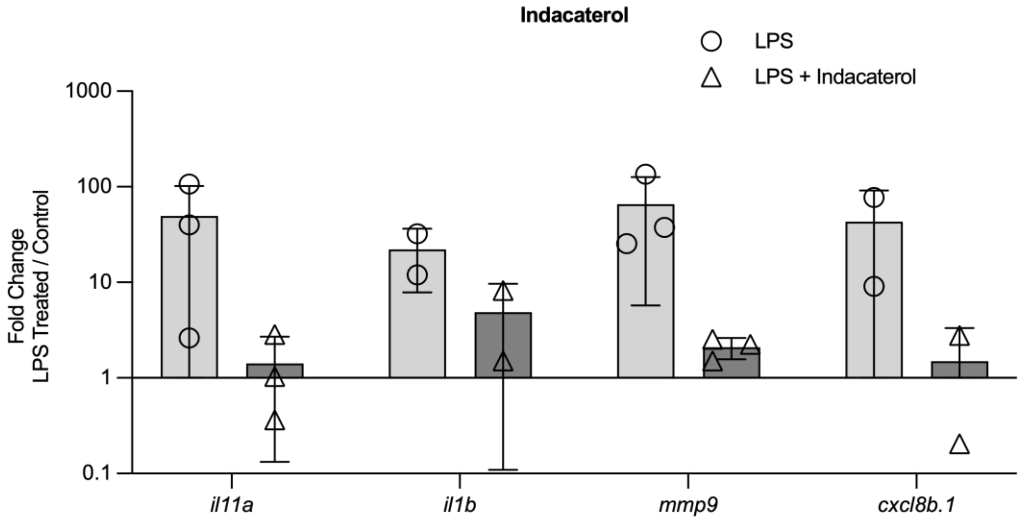


E

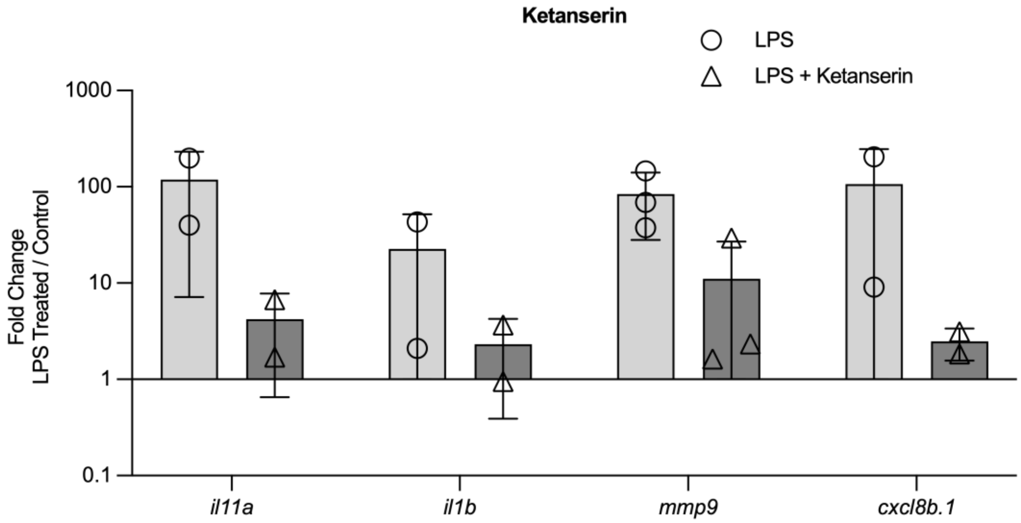


F

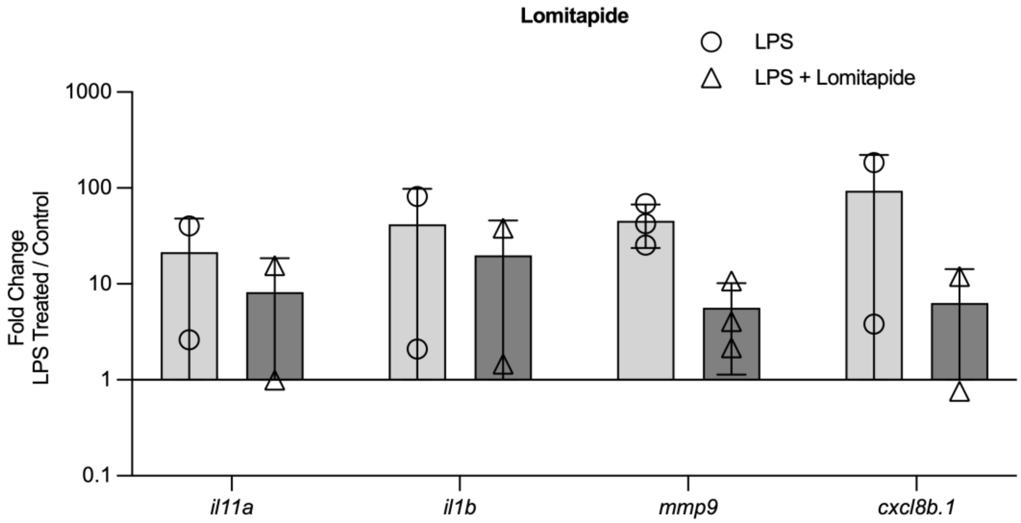

#### Figure S2

G

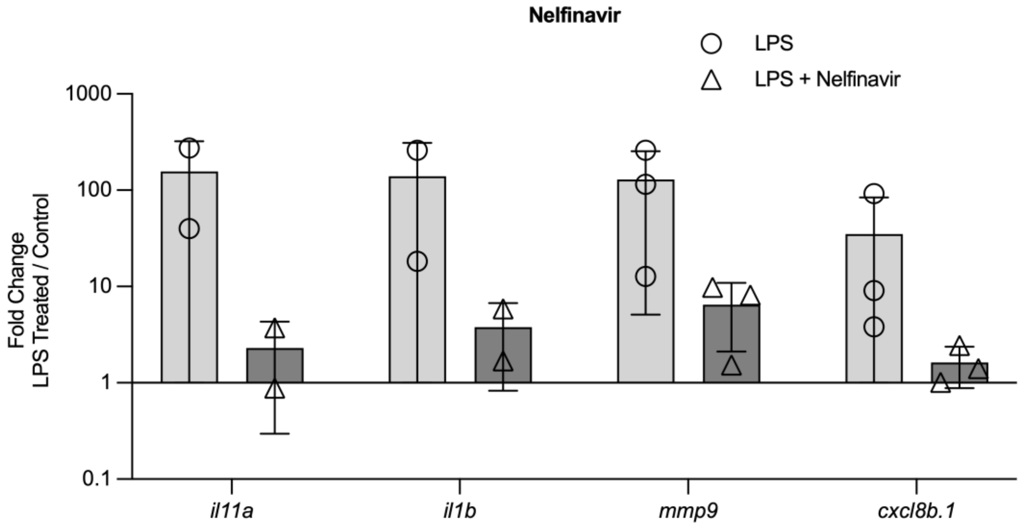


H

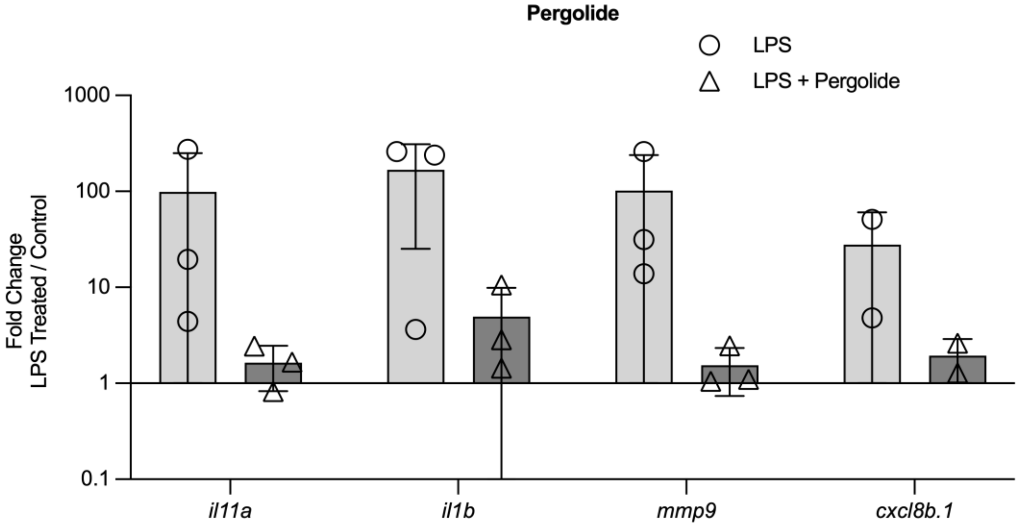


I

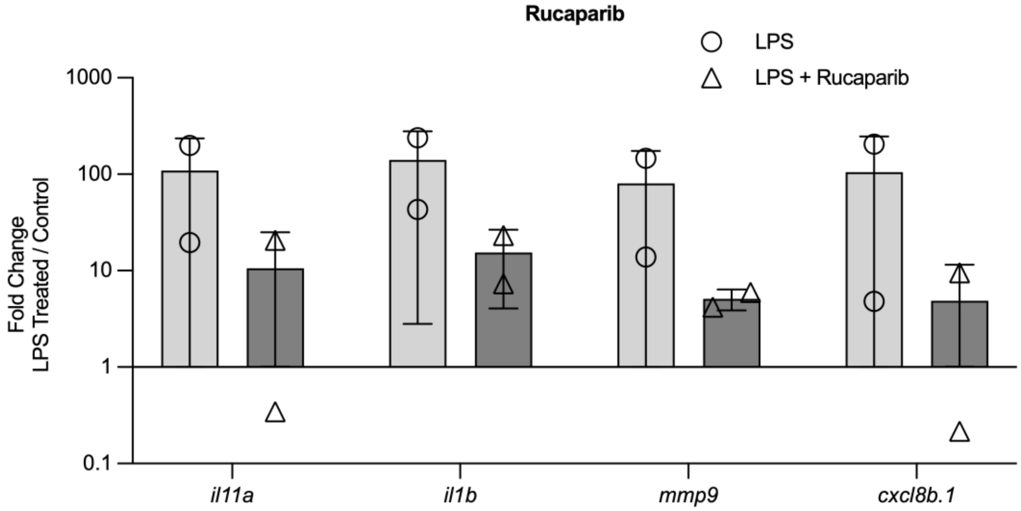

**Figure S2**

J

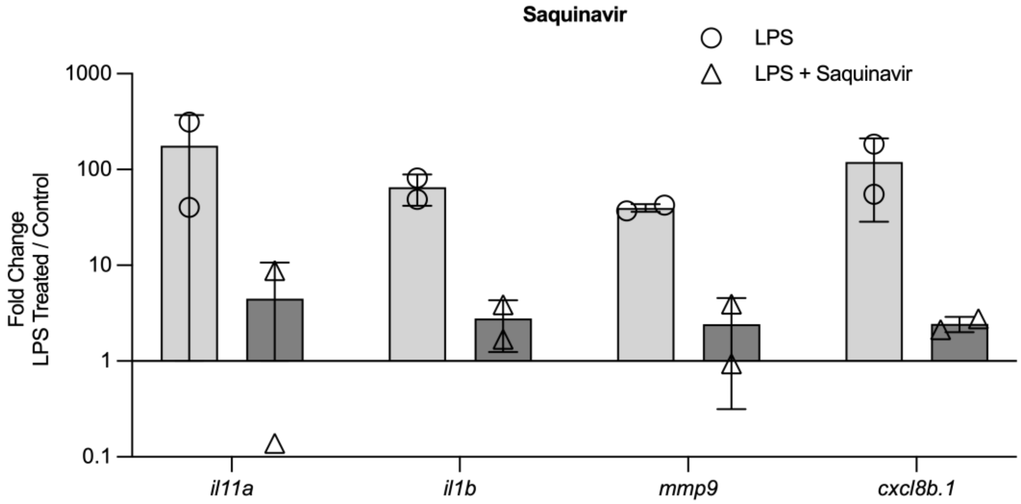


K

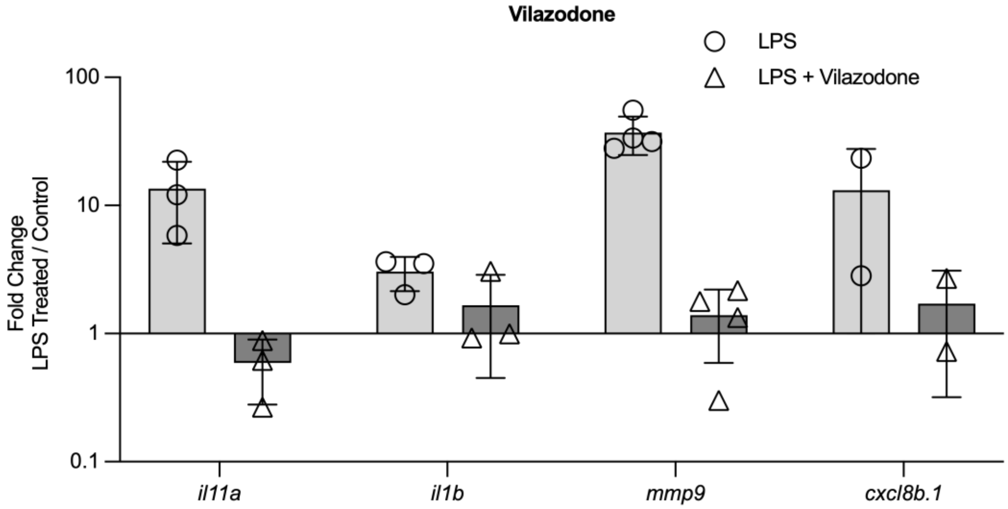


L

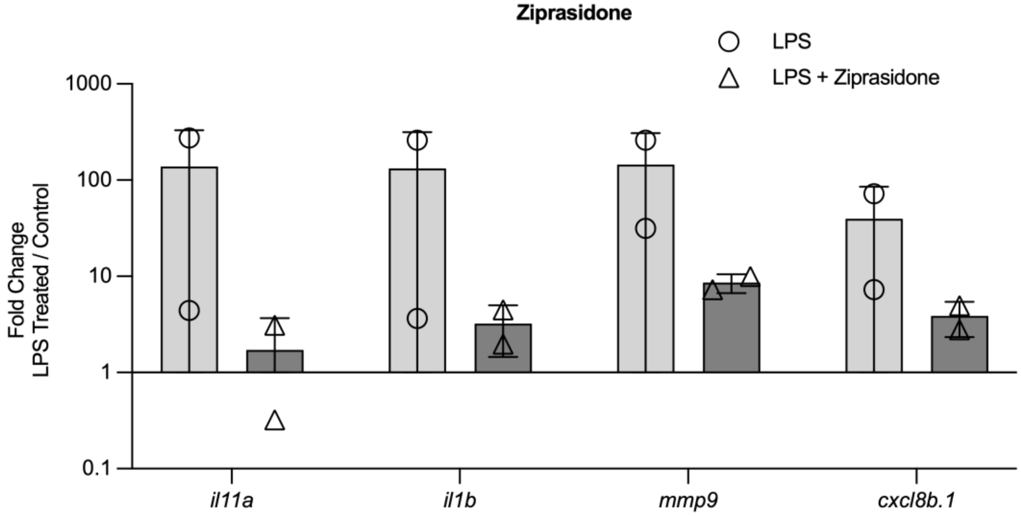

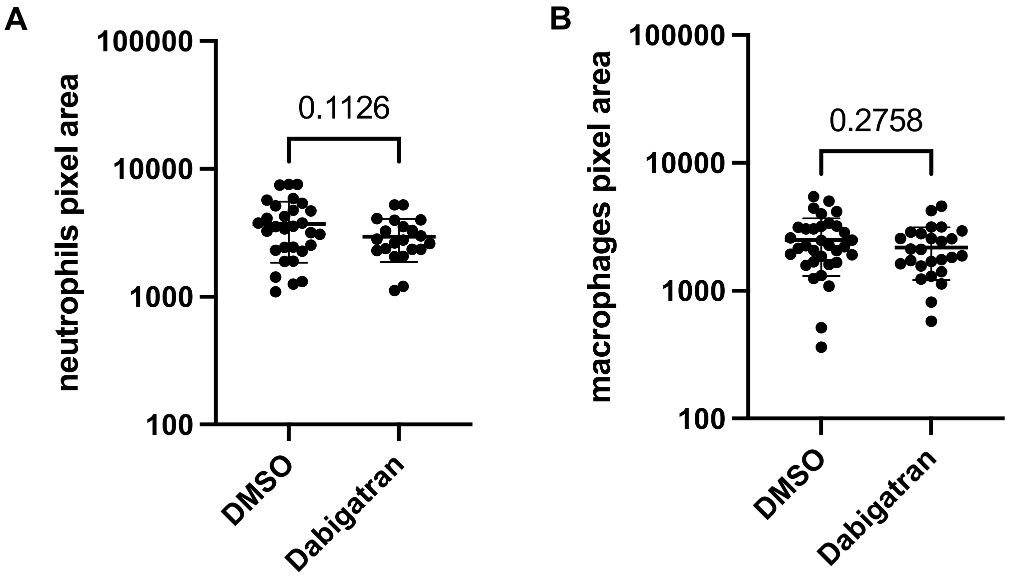

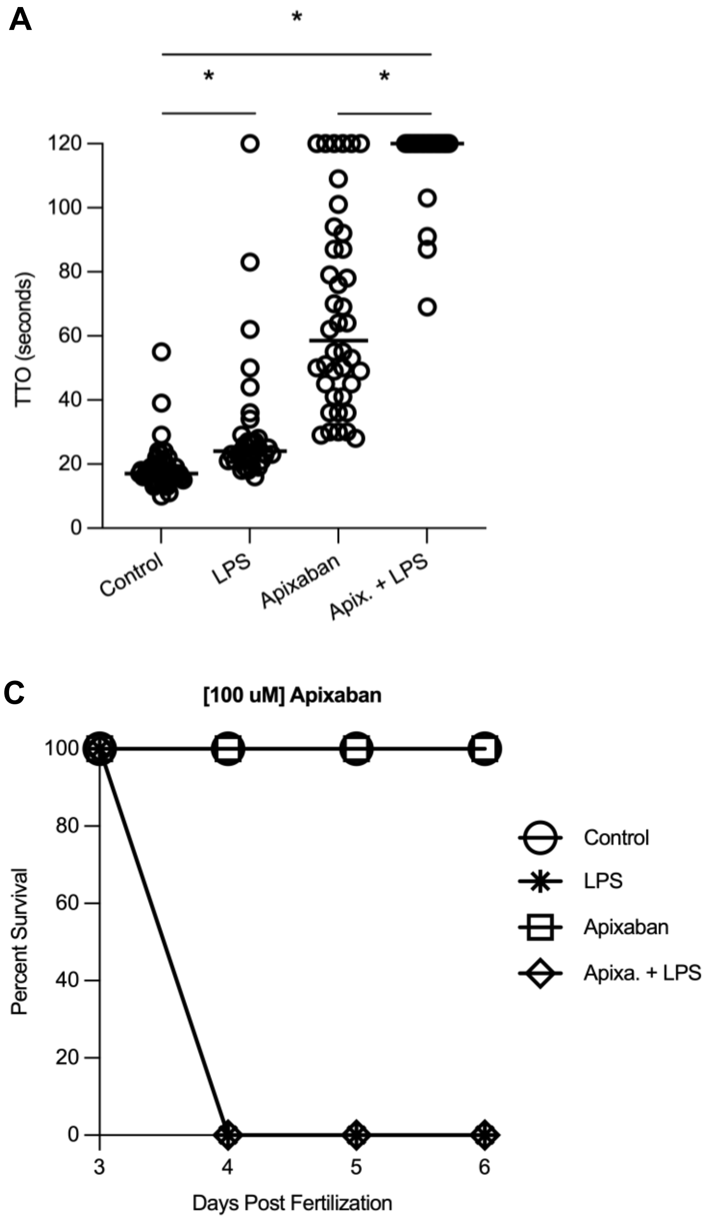
