## Supplemental Figure Legends for "Dabigatran prevents lipopolysaccharide mediated apoptosis in zebrafish through a thrombin independent mechanism"

**Figure S1: Multiple FDA-approved drugs protect zebrafish larvae from a lethal dose of LPS.** (A)-(U) Zebrafish were administered a lethal dose of LPS along with compounds from an FDA-approved repurposing library, and survival was tracked daily from 3 through 6 dpf. Any experimental controls comprised the same solvent volume of each drug, either DMSO or water.

**Figure S2: Co-administration of identified drugs with LPS decreases inflammatory cytokine expression.** (A)–(L) Expression of inflammatory genes was evaluated by qPCR 3 hours after drug administration in 3 dpf larvae treated with LPS, and each represented small molecule. Pooled larvae (15-45) from a single clutch were used, where each point represents a separate clutch, and the average fold change in LPS-treated fish vs. controls is graphed. At least two biological replicates were performed. Bars indicate mean and +/- 1 standard deviation. Student’s t-test on ΔΔC_t_ values was used to determine significance. *P<0.05.

**Figure S3: Tail transection does not result in increased neutrophil or macrophage migration to the site of injury.** 5-day-post-fertilisation (dpf) Tg(lyzC:GFP)^nz117^ and TgBAC (mpeg1.1: EGFP)^vcc7^ transgenic embryos were anesthetized and tail transaction was performed using a sterile surgical blade at the distal end of the notochord. After transection, embryos were transferred into fresh E3 medium containing 25 μM dabigatran and kept at 28.5 °C. The recruitment of neutrophils (A) and macrophages (B) to the wound site was imaged at 5 hours post transection and quanitified by measuring fluorescence at the wound site.

**Figure S4: Other direct anticoagulants do not protect zebrafish larvae from LPS toxicity.** (A),(B). Time to occlusion (TTO) after laser injury in the venous circulation was measured in 3 dpf zebrafish larvae 3 hours after LPS and anticoagulant administration. Each point represents a single larva, and the observer was blinded to the condition. P<0.05 was considered significant by Mann-Whitney *U* testing. (C), (D) Zebrafish were treated with apixaban or rivaroxaban and/or LPS at 3 dpf and observed daily until 6 dpf for survival.
